# Fibrillar α-synuclein induces neurotoxic astrocyte activation via RIP kinase signaling and NF-κB

**DOI:** 10.1101/2020.11.17.387175

**Authors:** Tsui-Wen Chou, Nydia P Chang, Medha Krishnagiri, Aisha P Patel, Colm Atkins, Brian P. Daniels

## Abstract

Parkinson’s disease (PD) is a neurodegenerative disorder characterized by death of midbrain dopamine neurons. The pathogenesis of PD is poorly understood, though misfolded and/or aggregated forms of the protein α-synuclein have been implicated in several neurodegenerative disease processes, including neuroinflammation and astrocyte activation. Astrocytes in the midbrain play complex roles during PD, initiating both harmful and protective processes that vary over the course of disease. However, despite their significant regulatory roles during neurodegeneration, the cellular and molecular mechanisms that promote pathogenic astrocyte activity remain mysterious. Here, we show that α-synuclein preformed fibrils (PFFs) induce pathogenic activation of human midbrain astrocytes, marked by inflammatory transcriptional responses, downregulation of phagocytic function, and conferral of neurotoxic activity. These effects required the necroptotic kinases RIPK1 and RIPK3, but were independent of MLKL and necroptosis. Instead, both transcriptional and functional markers of astrocyte activation occurred via RIPK-dependent activation of NF-κB signaling. Our study identifies a previously unknown function for α-synuclein in promoting neurotoxic astrocyte activation, as well as new cell death-independent roles for RIP kinase signaling in the regulation of glial cell biology and neuroinflammation. Together, these findings highlight previously unappreciated molecular mechanisms of pathologic astrocyte activation and neuronal cell death with implications for Parkinsonian neurodegeneration.

## Introduction

Parkinson’s disease (PD) is the second most common neurodegenerative disease after Alzheimer’s disease (1). Pathologically, PD is characterized by progressive loss of dopaminergic neurons in the substantia nigra pars compacta (SNpc), as well as axonal degeneration in the nigrostriatal pathway and reduction of dopamine inputs into the striatum (2). Though the pathogenesis of PD is poorly understood, growing evidence implicates aggregated forms of the protein α-synuclein as an etiologic agent of Parkinsonian neurodegeneration (3, 4). While normally serving as a lipid-binding protein which functions in vesicular trafficking, misfolding of soluble α-synuclein monomers leads to formation of insoluble aggregates (5). These aggregates exert neurotoxic and inflammatory activity which are thought to underlie the cell death and degeneration observed in PD and other synucleinopathies (6, 7).

While the impact of aggregated α-synuclein on neurons has been extensively described, roles for α-synuclein in astrocytes are comparatively poorly understood. During homeostasis, astrocytes serve key functions such as promoting neurite outgrowth and synapse formation, as well as phagocytosing cellular debris (8). However, the activation and proliferation of astrocytes is a hallmark of neuroinflammatory diseases, during which activated astrocytes have been ascribed both protective and pathologic functions (9-12). Reactive astrocytes are detectable in many neurodegenerative diseases, including PD (13, 14), Alzheimer’s disease (15), Huntington’s disease (16), amyotrophic lateral sclerosis (17, 18), and multiple sclerosis (19, 20). Upon activation, reactive astrocytes can exert both neuroinflammatory and neurotrophic effects (8, 19-23). Previous work has described at least two putative subtypes of reactive astrocytes, termed “A1,” which are proinflammatory and neurotoxic, and “A2,” which are generally anti-inflammatory and neurotrophic (19). These activation states have been distinguished by distinct transcriptional signatures and functional profiles, including the propensity to induce cell death in neurons (a marker of A1 astrocyte activity). While aggregated α-synuclein has been shown to contribute to inflammatory astrocyte activation (24-26), the transcriptional and functional consequences of this effect have not been fully established. Moreover, the molecular mechanisms that promote inflammatory astrocyte activation downstream of pathogenic α-synuclein species are poorly defined.

During neurodegeneration, inflammatory signals induce multiple forms of programmed cell death in susceptible neural cells, including both apoptosis and necroptosis (27-31). Necroptosis is a form of programmed cell death mediated by receptor-interacting protein kinases-1 (RIPK1) and -3 (RIPK3). These kinases coordinate activation of the executioner pseudokinase mixed lineage kinase domain-like protein (MLKL). Activated, oligomerized MLKL then permeabilizes the cell membrane, resulting in programmed necrotic cell death (32, 33). However, aside from necroptosis, several groups have recently described pleiotropic, cell death-independent functions for this pathway (34-40). We and others have shown that, in neurons, RIPK activation can initiate inflammatory transcriptional responses without inducing MLKL oligomerization and host cell death following neurotropic viral infection (34, 35, 41). However, whether necroptosis-independent functions for RIPK signaling are relevant during sterile neurodegenerative diseases requires further investigation. Moreover, whether RIPK signaling promotes inflammatory signaling in non-neuronal cells of the central nervous system (CNS), such as astrocytes, has yet to be fully addressed.

In this study, we sought to define the impact of aggregated α-synuclein on astrocyte activation state. We show that α-synuclein preformed fibrils (PFFs) induce robust inflammatory transcriptional signaling in human midbrain astrocytes, including transcripts associated with both the putative A1 and A2 astrocyte activation states. Functional analyses demonstrated that α-synuclein PFFs conferred neurotoxic activity in midbrain astrocytes, while diminishing homeostatic phagocytic activity. Each of these transcriptional and functional outcomes required the inflammatory transcription factor NF-κB, but not the JAK/STAT or AP1 pathways. Importantly, NF-κB activation and subsequent inflammatory transcription and neurotoxic activation could be rescued via pharmacological blockade of RIPK1 and RIPK3 signaling, while MLKL was dispensable for these effects, which occurred in the absence of astrocytic cell death. These data identify a previously unknown necroptosis-independent function for RIPK signaling in promoting a neurotoxic activation state in astrocytes following exposure to a pathogenic species of α-synuclein.

## Results

### α-synuclein PFFs induce transcriptional activity associated with astrocyte activation

To assess the impact of α-synuclein aggregates on astrocyte activation state, we treated primary cultures of human midbrain astrocytes with α-synuclein PFFs, which have been extensively shown to induce inflammatory activation and seed aggregation of endogenous α-synuclein both *in vitro* and *in vivo* (42-44). To profile astrocyte activation state following this treatment, we performed a qPCR screen of transcripts previously associated with the putative A1 (neurotoxic) and A2 (neurotrophic) astrocyte activation states (19), along with other general markers of inflammatory activity. To further characterize the nature of the PFF-induced transcriptional response, we performed these experiments in the presence of inhibitors of three major inflammatory transcriptional pathways: BAY 11-7085 (BAY) is an irreversible inhibitor of IκB Kinase (IKK), thereby blocking NF-κB activation; Pyridone 6 (PYR) is a pan-JAK inhibitor, thereby blocking signaling through STAT family transcription factors; and SR 11302 (SR) is an inhibitor of the transcription factor AP1.

Overnight treatment with α-synuclein PFFs induced increased expression of 8 of 11 A1-associated transcripts in our screen **(Figure 1a-g, Supplemental Figure 1a-b)**, including *HLA-A, SERPING1, SRGN, HLA-E, PSMB8, GBP2, FKBP5*, and *UGT1A1*, while expression of *GGTA1, FBLN5*, and *AMIGO2* was not significantly impacted by PFF treatment **(Supplemental Figure 1c-e)**. Notably, inhibition of NF-κB signaling with BAY rescued the upregulation of all 8 A1-associated transcripts, while inhibition of JAK/STAT or AP1 signaling had minimal effect. Surprisingly, however, PFF treatment also upregulated 5 of 11 A2-associated genes in our study (**(Figure 1h-m)**, including *PTX3, PTGS2, SPHK1, TM4FS1*, and *CLCF1*, while *S100A10, SLC10A6, B3GNT5, CD14, CD109*, and *EMP1* each showed various degrees of upregulation that did not reach statistical significance **(Figure 1n, Supplemental Figure 1f-j)**. PFF treatment also increased expression of a number of key neuroinflammatory chemokines, including *CCL2, CXCL1, CXCL10* **(Figure 1o-q)**. The upregulation of CXCL10, in particular, was also detected at the protein level via ELISA **(Figure 1r)**. In all cases, blockade of NF-κB signaling, but not JAK/STAT or AP1 signaling, prevented upregulation of reactive astrocyte genes. Together, these data suggest that α-synuclein PFFs strongly induce an NF-κB-dependent transcriptional response that includes a broad array of genes associated with astrocyte activation and inflammatory activity. However, the gene expression profile we observed did not fit neatly within the A1/A2 paradigm.

**Figure 1.**
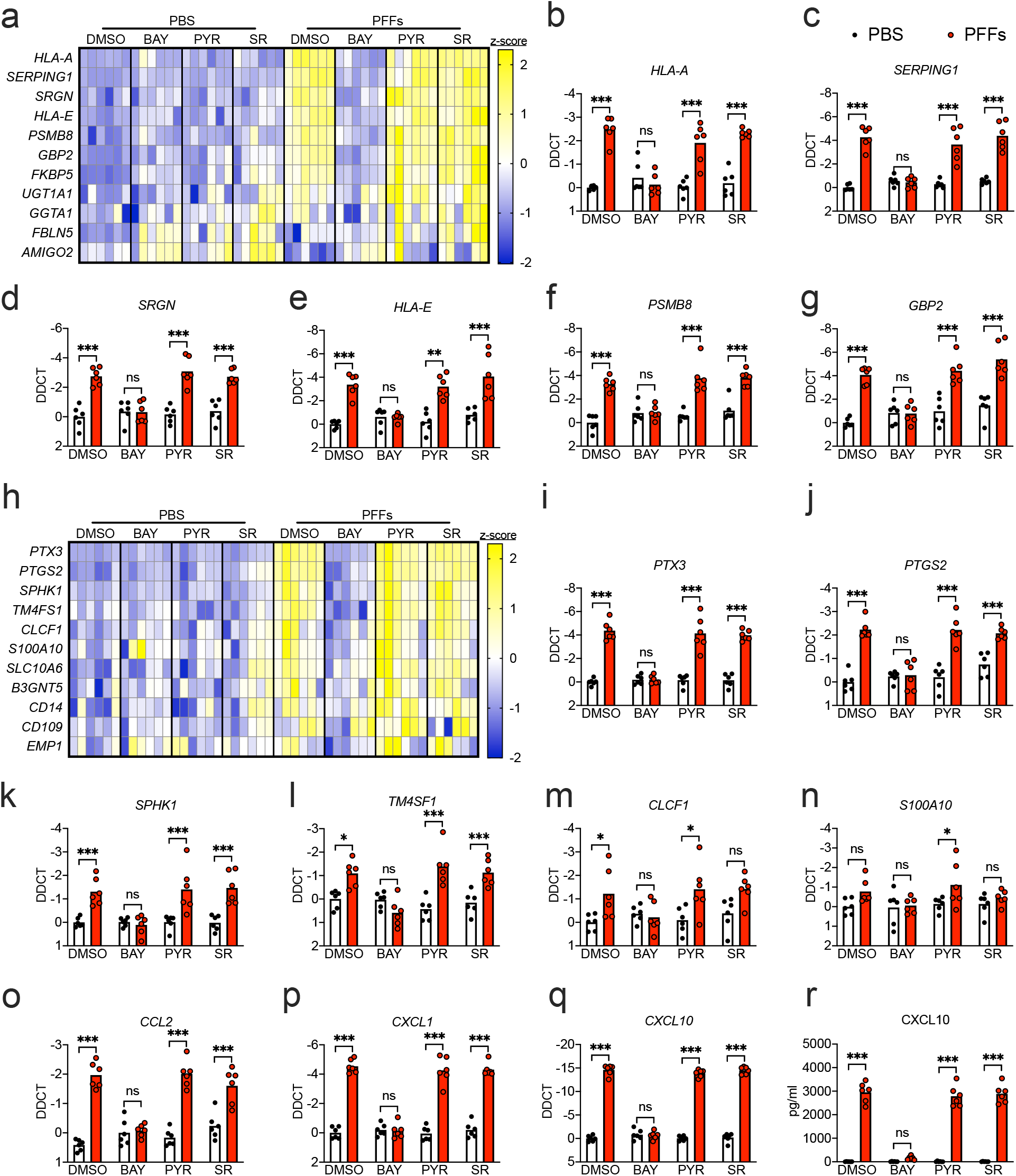
α-synuclein PFFs induce NF-κB-dependent transcriptional activation in human midbrain astrocytes. **a-r)** Primary human midbrain astrocyte cultures were treated for 24h with PFFs or PBS control solution. Cultures were pretreated (30min) with inhibitors of NF-κB (BAY), JAK/STAT (PYR), or AP1 (SR) signaling prior to addition of PFFs. Levels of indicated A1-associated transcripts **(a-g)**, A2-associated transcripts **(h-n)**, and inflammatory chemokines **(o-q)** were measured using qRT-PCR. **r)** Levels of CXCL10 protein in culture supernatants were measured via ELISA. Data in **(a)** and **(h)** are normalized and z-transformed summaries of the qPCR data that appears in the bar graphs throughout the figure and Supplemental Figure 1, represented via heatmaps. ns: not significant, *p < 0.05, **p < 0.01, ***p < 0.001. Bars represent group means. n=6 independent replicates for all experiments.

### α-synuclein PFFs induce both expression and activation of NF-κB signaling elements

We next sought to more thoroughly assess the impact of α-synuclein PFFs on NF-κB activation in astrocytes. While BAY blocks NF-κB signaling by inhibiting upstream IKK activity, we tested whether direct blockade of NF-κB nuclear translocation using the inhibitor JSH-23 would also inhibit inflammatory transcription in PFF-treated astrocytes. We, indeed, observed that JSH-23 treatment rescued the induction of several reactive astrocyte genes following PFF treatment, including *SERPING1, HLA-E, GBP2*, and *CXCL10* **(Figure 2a-d)**. To directly confirm that PFFs induced NF-κB activation in astrocytes, we performed confocal microscopy to visualize nuclear translocation of the NF-κB component p65. PFF treatment induced robust accumulation of p65 in the nucleus, as indicated by enhanced colocalization of p65 signal with nuclear DAPI staining **(Figure 2e-f)**. Importantly, treatment with BAY completely blocked this effect, confirming that IKK inhibition effectively blocked NF-κB activation in our experiments. As a secondary confirmation of NF-κB activation, we extracted nuclear fractions from astrocytes following treatment with PFFs or PBS control and performed ELISA to detect p65. These experiments also revealed significant accumulation of nuclear p65 which was completely blocked by cotreatment with BAY **(Figure 2g)**.

**Figure 2.**
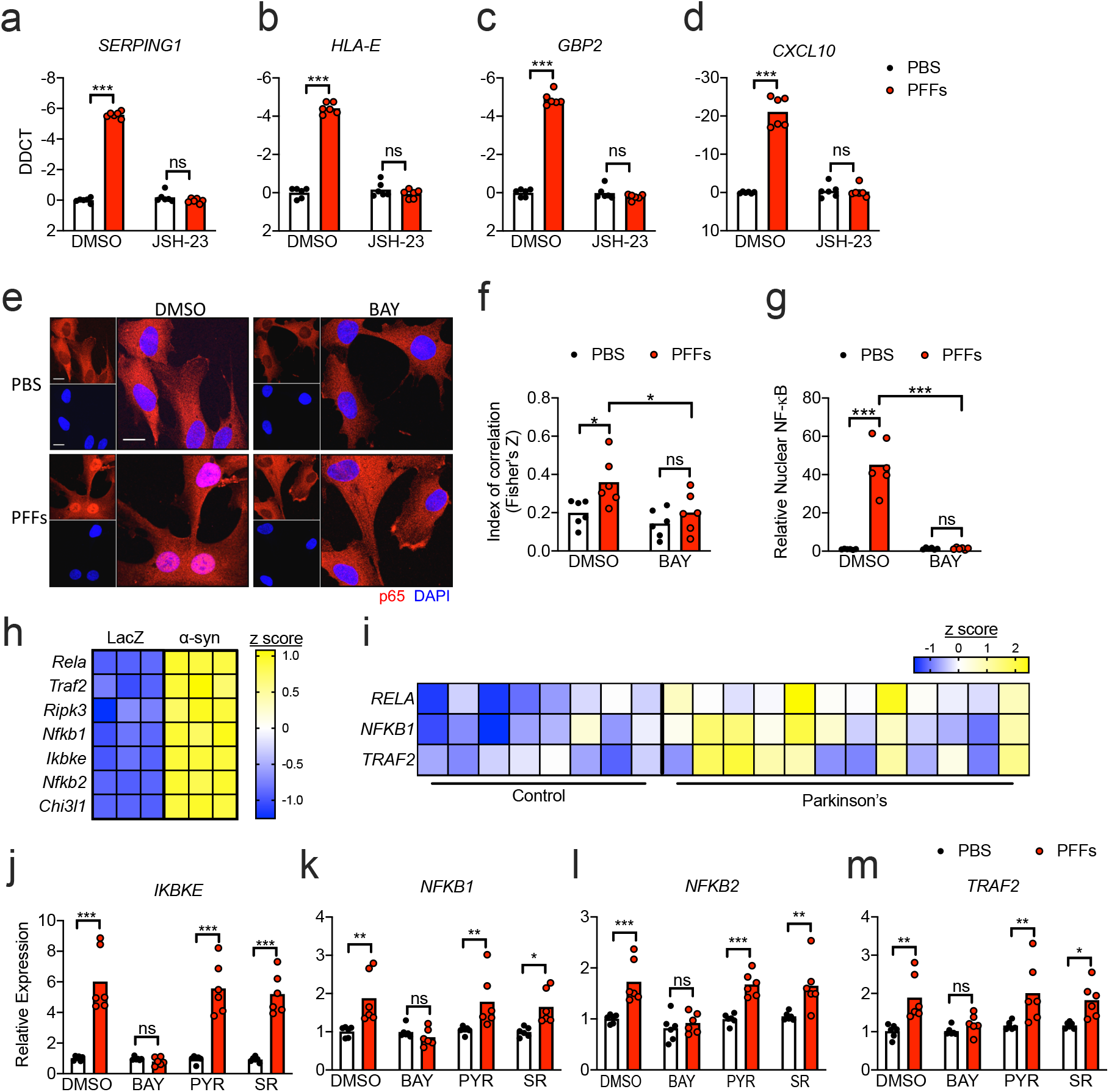
α-synuclein induces NF-kB expression and activation in astrocytes. **a-d)** qRT-PCR analysis of indicated genes in human midbrain astrocyte cultures treated following 24h PFF treatment +/- cotreatment with JSH-23. **e)** Confocal microscopy of NF-κB component p65 (red) and nuclei (DAPI, blue) in human midbrain astrocytes following 2h PFF treatment +/- cotreatment with BAY. Scale bars = 20 µm. **f)** Colocalization of red and blue signal in (e) is quantified as Fisher’s Z transformed index of correlation. **g)** Nuclear fractions were isolated from human midbrain astrocyte cultures following 2h PFF treatment +/- cotreatment with BAY. Relative levels of p65 in nuclear fractions were quantified using ELISA. **h-i)** Secondary analysis of microarray profiling of **h)** rat astrocytes treated with conditioned neuronal medium containing LacZ or α-synuclein (GSE11574) or **i)** human postmortem substantia nigra samples from patients with PD or healthy controls (GSE26927). Z scores were calculated for genes under the GO term 0038061 and those exhibiting significantly differential expression between groups (corrected p value <0.05) are displayed in the respective heatmaps. **(j-m)** qRT-PCR analysis of indicated genes in human midbrain astrocyte cultures treated following 24h PFF treatment +/- cotreatment with indicated inhibitors. ns: not significant, *p < 0.05, **p < 0.01, ***p < 0.001. Bars represent group means. n=6 independent replicates for all experiments.

We also questioned whether PFF treatment might influence NF-κB activity via upregulation of NF-κB components and signaling elements. To answer this, we first turned to two publicly available transcriptomic databases. In one study, Lee and colleagues (25) exposed primary rat astrocytes to conditioned medium from the human neuroblastoma cell line SH-SY5Y that had been transduced to express either α-synuclein or LacZ control. Secondary analysis of microarray data from this study revealed that exposure to α-synuclein-containing conditioned medium significantly upregulated many genes associated with the “NF-κB signaling” gene ontology term (GO:0038061), including *Rela* (p65), *Traf2, Ripk3, Nfkb1, Ikbke, Nfkb2*, and *Chi3l1* **(Figure 2h)**. We performed a similar analysis on a dataset originally published by Durrenberger and colleagues (45), who reported transcriptomic profiles of postmortem samples of substantia nigra tissues from 12 PD patients and 8 healthy controls. Secondary analysis of this dataset showed significant upregulation of 3 genes associated with NF-κB signaling, including *RELA, NFKB1*, and *TRAF2* **(Figure 2i)**. In our own experimental system, we saw that treatment of human midbrain astrocytes with α-synuclein PFFs significantly upregulated expression of *IKBKE, NFKB1, NFKB2*, and *TRAF2* **(Figure 2j-m)**. Notably, the induction of these genes was blocked by BAY, suggesting that PFFs may induce feed-forward amplification of NF-κB signaling in astrocytes. Together, these data confirm that α-synuclein induces both expression and activation of NF-κB in astrocytes and that enhanced expression of NF-κB signaling elements in the substantia nigra is a feature of PD in human patients.

### α-synuclein PFFs induce neurotoxic activity in midbrain astrocyte cultures

Beyond transcriptional activity, a key feature of astrocyte activation are changes to homeostatic function. A1, or neuroinflammatory astrocytes, generally, have been reported to induce programmed cell death in neurons, thereby contributing to disease pathogenesis. To test if α-synuclein PFFs could confer neurotoxic activity to astrocytes, we treated midbrain astrocyte cultures with PFFs or PBS control in the presence of transcription factor pathway inhibitors **(Figure 3a)**. We then collected astrocyte conditioned medium (ACM) from these cultures and added it to cultures of differentiated SH-SY5Y neuroblastoma cells at a 1:1 ratio with normal culture medium. We observed that 24h treatment with ACM derived from astrocyte cultures treated with PFFs significantly reduced the viability of SH-SY5Y cultures, as assessed via ATP luciferase assay **(Figure 3b)**. Blockade of NF-κB signaling in astrocytes with BAY rescued this neurotoxic activity, while PYR and SR treatment did not. We saw similarly that NF-κB inhibition with JSH-23 also prevented neurotoxic activity in astrocytes following PFF treatment **(Figure 3c)**. As a secondary confirmation of cell death in SH-SY5Y cultures, we performed terminal deoxynucleuotidyl transferase dUTP nick end labeling (TUNEL) to detect DNA damage associated with programmed cell death. This assay revealed a significant increase in the number of TUNEL^+^ nuclei in SH-SY5Y cultures treated with ACM derived from PFF-treated astrocytes **(Figure 3d-e)**. However, treatment of astrocytes with BAY prevented this effect. Together, these data suggest that α-synuclein PFFs stimulate neurotoxic activation in midbrain astrocytes that is dependent on NF-κB. Importantly, we confirmed that cell death in our study could not be explained by exposure to residual amounts of inhibitors found in ACM samples, as direct treatment of SH-SY5Y cultures with each inhibitor alone did not impact their viability (**Supplemental Figure 2**).

**Figure 3.**
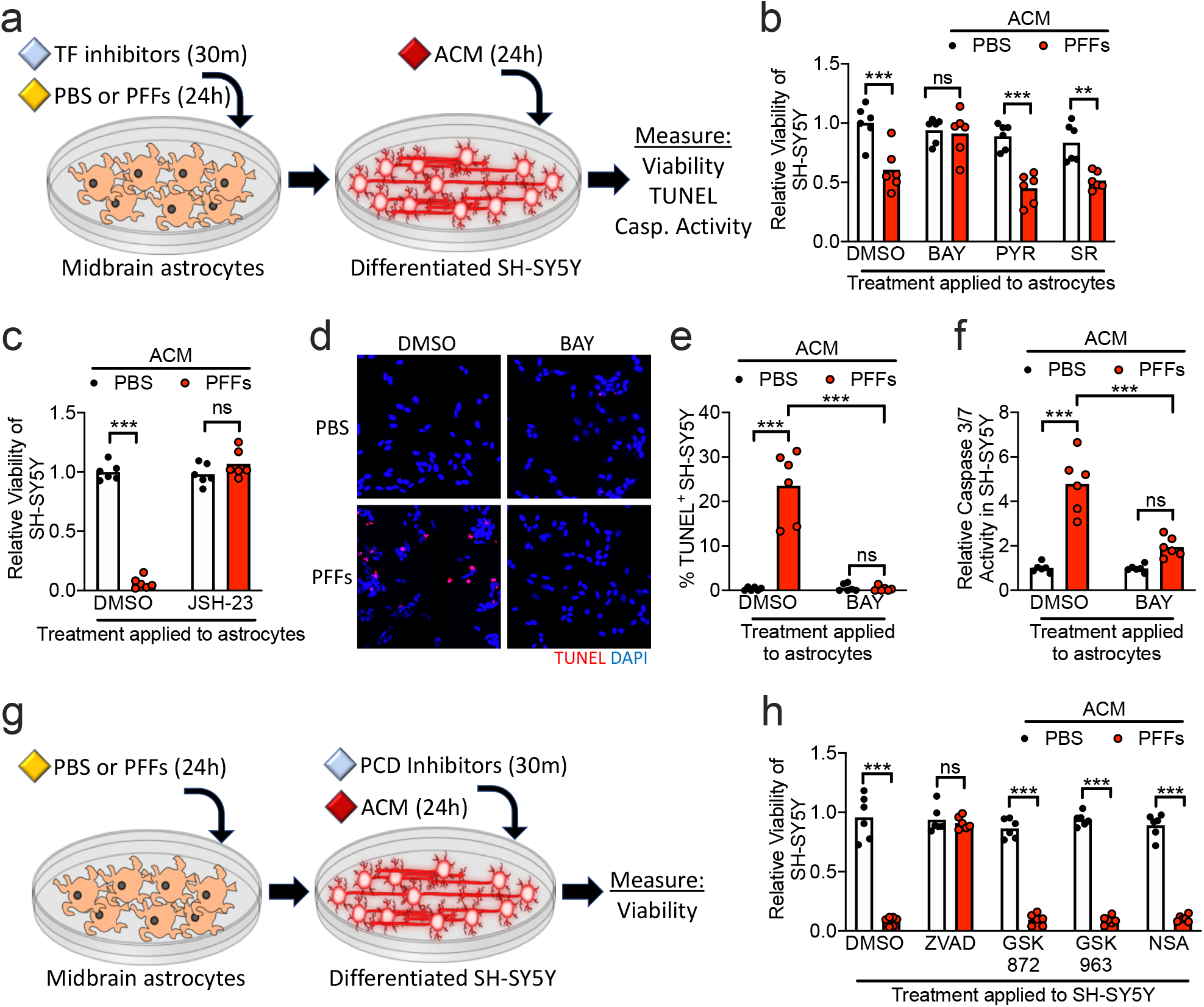
α-synuclein PFFs induce neurotoxic activity in midbrain astrocytes. **a-f)** Primary human midbrain astrocytes were treated with transcription factor (TF) inhibitors and/or PFFs, as indicated. After 24h, astrocyte conditioned medium (ACM) was applied (1:1) to differentiated SH-SY5Y cultures for 24h followed by endpoint analyses. **b-c)** Viability of SH-SY5Y cells following treatment with ACM derived from astrocyte cultures treated with the indicated inhibitors was measured via ATP-luciferase assay (Cell Titer Glo). **d-e)** Cell death of SH-SY5Y cells following 24h ACM treatment was detected via TUNEL **(d)** and quantified as percentage of TUNEL^+^ nuclei **(e). f)** Levels of Caspase 3/7 activity in SH-SY5Y cells following treatments as in **(a)** was measured using a chromogenic DEVD-cleavage assay. **g)** Modification of experimental setup in **(a)**, in which SH-SY5Y cultures were pretreated with programmed cell death (PCD) inhibitors for 30m prior to addition of ACM. **h)** Viability of SH-SY5Y cells following treatment with ACM and PCD inhibitors as in **(g)** was measured via ATP-luciferase assay (Cell Titer Glo). ns: not significant, **p<0.01, ***p < 0.001. Bars represent group means. n=6 independent replicates for all experiments.

While apoptosis is the most commonly reported form of cell death in the brain during Parkinsonian neurodegeneration, contributions of other cell death modalities, including necroptosis, have also been reported (28, 46). We questioned which cell death modality was induced in differentiated SH-SY5Ycells in our experiments. We first used a chromogenic DEVD-cleavage assay to measure levels of executioner caspase (caspase 3 and caspase 7) activity in ACM-treated SH-SY5Y cultures. These experiments revealed that ACM derived from PFF-treated astrocytes induced robust Caspase 3/7 activity in SH-SY5Y cultures, and this effect was blocked when astrocytes were cotreated with BAY **(Figure 3f)**. To more carefully determine if caspase-dependent apoptosis was occurring, we performed additional experiments in which we modified our treatment paradigm. Prior to addition of ACM, SH-SY5Y cultures were pretreated with inhibitors of programmed cell death, including the pan-caspase inhibitor zVAD-FMK (zVAD), the RIPK3 inhibitor GSK872, the RIPK1 inhibitor GSK963, or the MLKL inhibitor necrosulfanamide (NSA) **(Figure 3g)**. Blockade of caspase signaling with zVAD completely rescued cell death observed in SH-SY5Y cultures following treatment with ACM derived from PFF-treated astrocytes, while inhibition of necroptosis signaling components (RIPK1, RIPK3, or MLKL) did not **(Figure 3h)**. These data suggest that the neurotoxic activity conferred by PFFs in astrocytes induces apoptosis rather than necroptosis in SH-SY5Y cells.

### α-synuclein PFFs reduce homeostatic phagocytic activity in astrocytes

Previous work has shown that neurotoxic activity in astrocytes occurs concomitantly with the downregulation of key homeostatic functions, including phagocytosis. We thus questioned whether α-synuclein PFFs would perturb the phagocytic function of midbrain astrocytes. We treated astrocyte cultures with PFFs or PBS control solution for 24h in the presence of transcription factor pathway inhibitors and performed qRT-PCR analysis of key genes known to be involved in astrocytic phagocytosis. PFF treatment downregulated expression of *GAS6* **(Figure 4a)**, which encodes an important opsonin for phagocytosis of apoptotic cells, as well as *MEGF10* **(Figure 4b)**, which encodes a receptor for C1q, an important opsonin in the classical complement pathway. In contrast, expression of *MERTK*, a GAS6 receptor, was increased by PFF treatment **(Figure 4c)**, which may represent a compensatory response to decreased GAS6 expression. Expression of *AXL*, which encodes an additional GAS6 receptor, was not impacted **(Figure 4d)**. BAY treatment prevented all PFF-induced changes to phagocytic gene expression, while PYR and SR had marginal or no impact. To determine whether changes to phagocytic gene expression actually impacted phagocytic activity, we measured uptake of fluorescently labelled zymosan, a yeast-associated glucan, in astrocyte cultures following 24 treatment with PFFs and/or BAY (or respective controls). PFF-treated astrocytes phagocytosed significantly less zymosan compared to control cultures, and this effect was blocked in the presence of BAY **(Figure 4e)**. These data suggest that, in addition to inducing inflammatory gene transcription and neurotoxic activity, NF-κB activation downstream of PFF treatment reduces the homeostatic phagocytic capacity of astrocytes.

**Figure 4.**
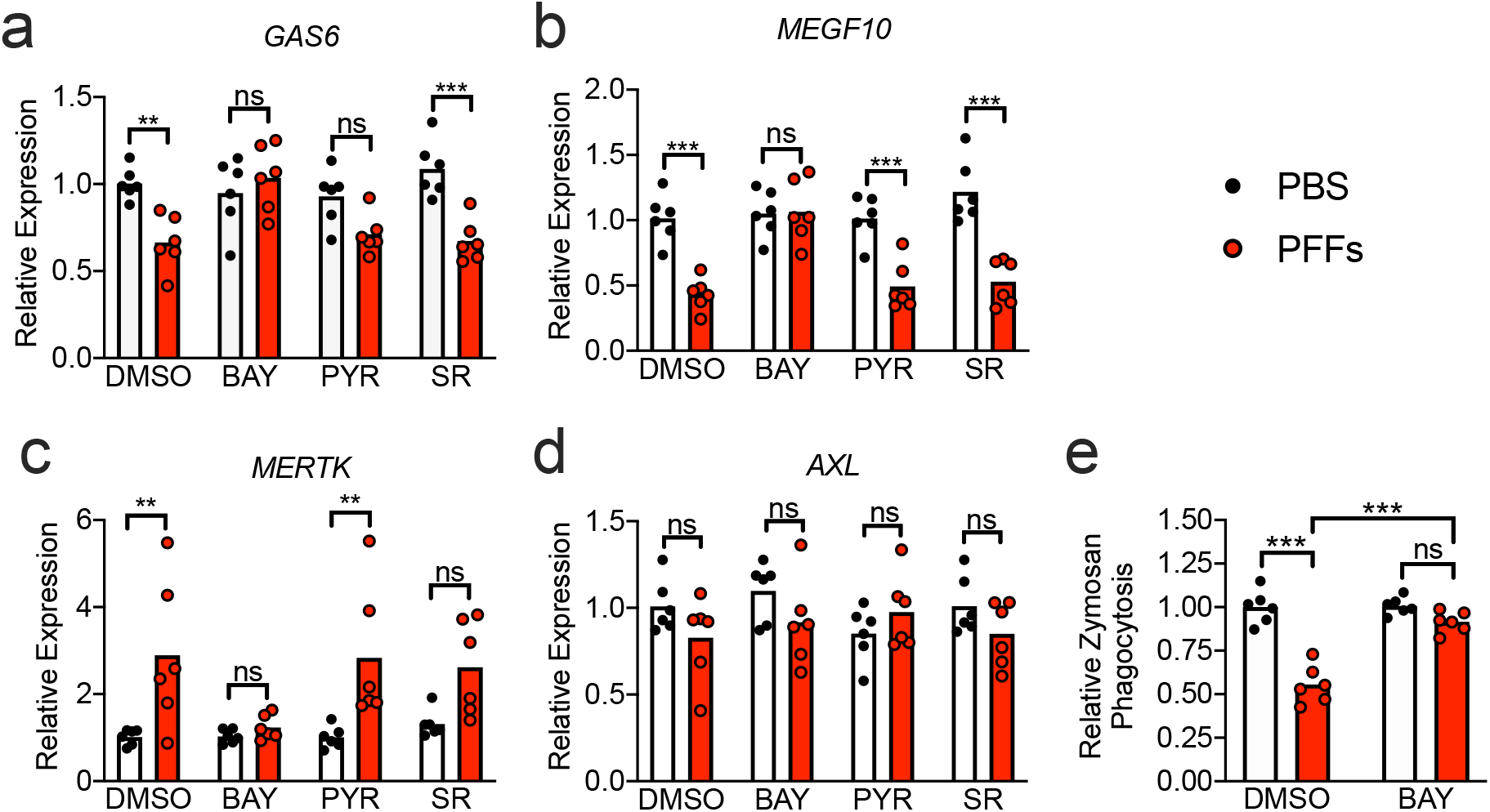
α-synuclein PFFs reduce phagocytic activity in midbrain astrocytes. **a-e)** Primary human midbrain astrocyte cultures were treated for 24h with PFFs or PBS control solution. Cultures were pretreated (30m) with inhibitors of NF-κB (BAY), JAK/STAT (PYR), or AP1 (SR) signaling prior to addition of PFFs. **a-d)** Levels of indicated phagocytosis-associated transcripts were measured using qRT-PCR. **e)** 24h following PFF or PBS treatment, relative rates of phagocytosis were assessed by measuring uptake of fluorescently labelled zymosan. ns: not significant, **p < 0.01, ***p < 0.001. Bars represent group means. n=6 independent replicates for all experiments.

### α-synuclein PFFs induce RIPK-dependent transcriptional activation in astrocytes, independently of necroptosis

While our data clearly identified NF-κB as a molecular driver of astrocyte activation in our system, the upstream inputs into this pathway remained unclear. We previously described a role for RIPK signaling in inflammatory transcriptional activation in in neurons that was independent of necroptotic cell death. Moreover, RIPK signaling is a known activator of the NF-κB pathway (47-50). We thus questioned whether RIPK signaling was required for PFF-mediated astrocyte activation. Treatment of midbrain astrocyte cultures with PFFs for 24h in the presence of inhibitors of either RIPK3 or RIPK1 revealed that both kinases were required for induction of the PFF-mediated transcriptional response, including genes associated with astrocyte activation (*GBP2, HLA-E*, and *SERPING1*) **(Figure 5a-c)** and inflammatory chemokines (*CCL2, CXCL1, CXCL10*) **(Figure 5d-f)**. Blockade of RIPK signaling also prevented induction of CXCL10 protein expression, as confirmed by ELISA **(Figure 5g)**. We also observed that inhibition of both RIPK3 and RIPK1 prevented transcriptional induction of NF-κB associated genes, including *IKBKE, NFKB1, NFKB2*, and *TRAF2* **(Figure 5h-k)**. Notably, these effects were independent of necroptosis, as inhibition of MLKL had no effect on PFF-mediated transcriptional activation for any class of genes measured in our study **(Supplemental Figure 3a-f, h-k)**, nor did it impact protein expression of CXCL10 **(Supplemental Figure 3g)**. Moreover, neither PFFs nor any of the inhibitors in our study induced detectable levels of cell death in astrocytes **(Supplemental Figure 4)**, further confirming that necroptosis was not the source of transcriptional activation downstream of astrocytic RIPK signaling. These data suggest that RIPK1 and RIPK3 engage inflammatory transcriptional activity in astrocytes following PFF treatment via an MLKL-independent mechanism.

**Figure 5.**
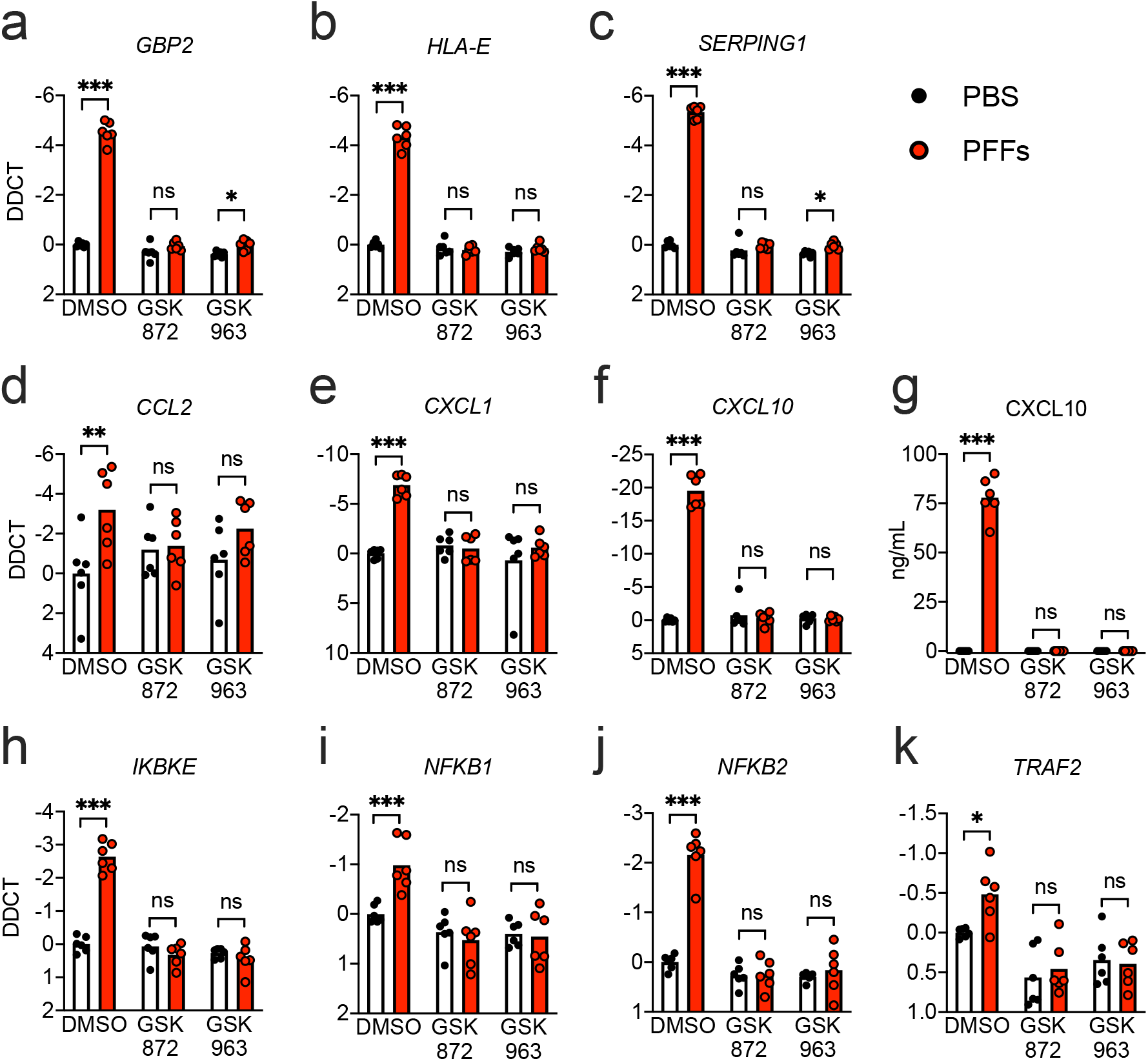
α-synuclein PFF-mediated transcriptional activation in astrocytes requires RIPK1 and RIPK3. **a-j)** Primary human midbrain astrocyte cultures were treated for 24h with PFFs or PBS control solution. Cultures were pretreated (30min) with inhibitors of RIPK3 (GSK872) or RIPK1 (GSK963) signaling prior to addition of PFFs. **a-f, h-k)** Levels of indicated transcripts were measured using qRT-PCR. **g)** Levels of CXCL10 protein in culture supernatants were measured via ELISA. ns: not significant, *p < 0.05, **p < 0.01, ***p < 0.001. Bars represent group means. n=6 independent replicates for all experiments.

### RIPK signaling is required for NF-κB activation and neurotoxic activity downstream of α-synuclein PFFs

We next confirmed that RIPK signaling was required for NF-κB activation in PFF-treated astrocytes. Confocal microscopic analysis of p65 expression revealed that blockade of either RIPK1 or RIPK3 was sufficient to prevent nuclear accumulation of p65, while blockade of MLKL had no effect on p65 nuclear translocation **(Figure 6a-b)**. We observed similar findings following detection of p65 in isolated nuclear fractions using ELISA **(Figure 6c)**. To confirm that RIPK signaling was likewise also required for functional indications of astrocyte activation, we treated midbrain astrocytes with PFFs or PBS control for 24h following 30m pretreatment with GSK872 or GSK963. We then treated differentiated cultures of SH-SY5Y cells with the ACM from these astrocyte cultures at a 1:1 ratio with normal culture medium **(Figure 6d)**. As expected, ACM derived from PFF-treated astrocytes greatly reduced the viability of SH-SY5Y cultures. However, this effect could be blocked by inhibition of either RIPK1 or RIPK3 in astrocytes **(Figure 6e)**. These data suggest that necroptosis-independent RIPK activity engages both transcriptional and functional activation of astrocytes following exposure to fibrillar α-synuclein. We thus returned to our secondary analysis of gene expression in the substantia nigra of Parkinson’s patients in order to see if there was evidence of increased expression of this pathway in human PD. We observed significant upregulation of *RIPK3* in PD patients compared to normal controls, while expression of both *RIPK1* and *MLKL* did not reach statistical significance **(Figure 6f)**. Together, these data identify a previously unknown function for the RIPKs in the promotion of a neurotoxic activation state and suggest further work is needed to identify roles for necroptosis-independent RIPK signaling in Parkinsonian neurodegeneration and other synucleinopathies.

**Figure 6.**
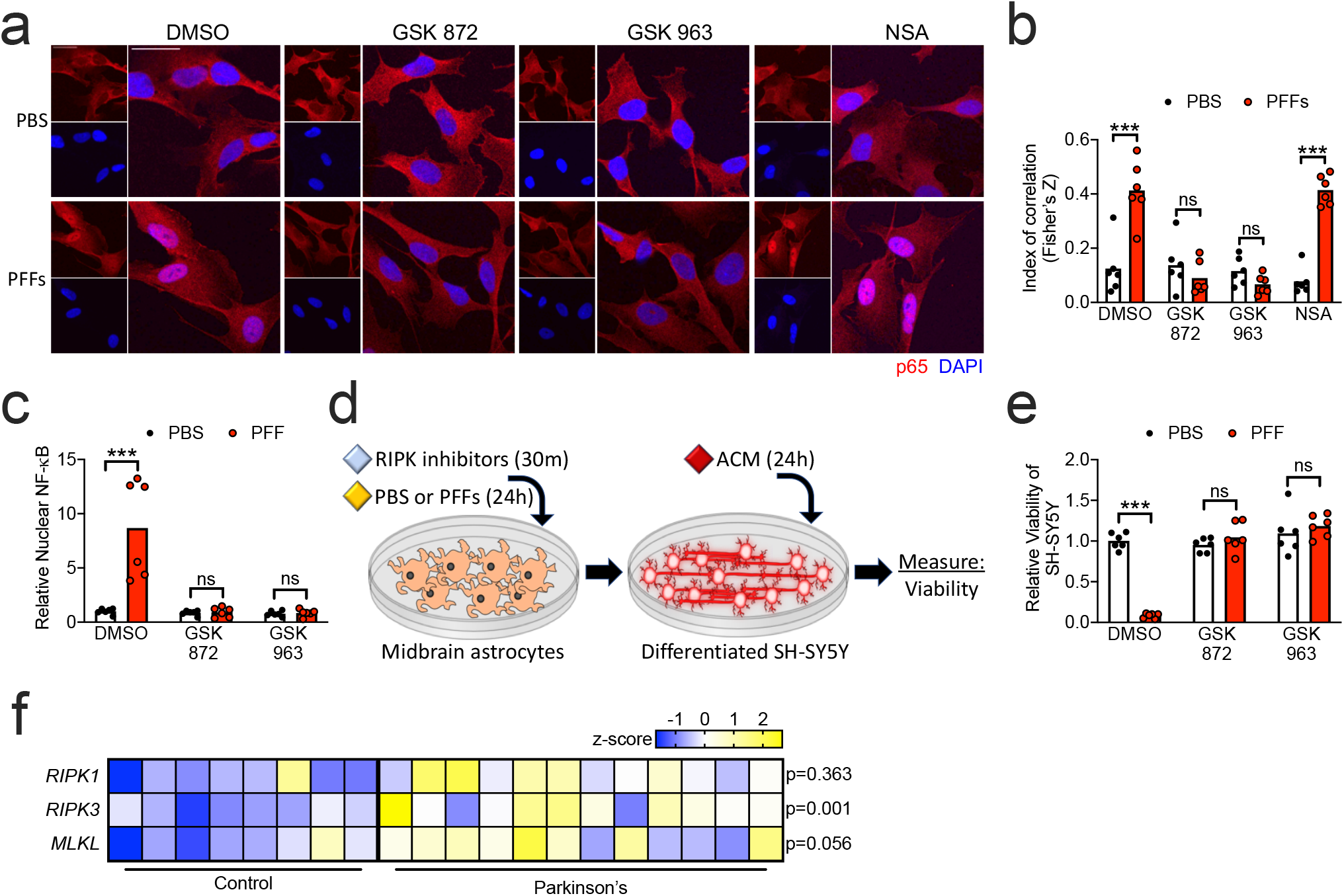
α-synuclein PFF-mediated transcriptional activation in astrocytes requires RIPK1 and RIPK3. **a)** Confocal microscopy of NF-κB component p65 (red) and nuclei (DAPI, blue) in human midbrain astrocytes following 2h PFF treatment +/- cotreatment with indicated inhibitors. Scale bars = 50 µm. **b)** Colocalization of red and blue signal in **(e)** is quantified as Fisher’s Z transformed index of correlation. **c)** Nuclear fractions were isolated from human midbrain astrocyte cultures following 2h PFF treatment +/- cotreatment with indicate inhibitors. Relative levels of p65 in nuclear fractions were quantified using ELISA. **d)** Primary human midbrain astrocytes were treated with RIPK inhibitors and/or PFFs, as indicated. After 24h, astrocyte conditioned medium (ACM) was applied (1:1) to differentiated SH-SY5Y cultures for 24h followed by endpoint analyses. **e)** Viability of SH-SY5Y cells following treatment with ACM derived from astrocyte cultures treated with the indicated inhibitors was measured via ATP-luciferase assay (Cell Titer Glo). **f)** Secondary analysis of microarray profiling of human postmortem substantia nigra samples from patients with PD or healthy controls (GSE26927). Z scores and corrected p-values were calculated for necroptosis pathway components indicated and displayed via heatmap. ns: not significant, *p < 0.05, **p < 0.01, ***p < 0.001. Bars represent group means. n=6 independent replicates for all experiments.

## Discussion

Misfolding and abnormal aggregation of α-synuclein is a well-known pathological hallmark of PD, during which it has been implicated in the formation of pathogenic Lewy bodies (LB) and dopaminergic neuron loss in the SNpc (51-53). However, the effects of pathogenic α-synuclein species on astrocytes have not been thoroughly studied. Previous work has shown that astrocytes can protect dopaminergic neurons from α-synuclein deposition and degeneration (54). However, α-synuclein aggregates are also frequently observed in PD patients, where they disturb vital homeostatic functions and thereby exacerbate disease pathology in neurons (55). While it is clear that aggregated α-synuclein induces inflammatory activity in astrocytes, the molecular mechanisms underlying this effect and their functional consequences in disease pathogenesis remain poorly understood. We show that α-synuclein PFFs induce a potently inflammatory transcriptional program in human midbrain astrocytes that is associated with neurotoxic activity and decreased homeostatic phagocytic function. Notably, this effect requires RIPK signaling, but does not result in necroptosis, the canonical function of this pathway. These data suggest a previously unknown function for RIPK signaling in the induction of a neurotoxic activation state in astrocytes following exposure to fibrillar α-synuclein. Notably, RIPK3 activation in astrocytes has been observed in mouse models of PD (29), as well as postmortem human patient samples (28). Moreover, both genetic and pharmacological ablation of RIPK3 signaling have been shown to be protective in mouse models of neurodegeneration (56, 57). Our findings suggest that therapies targeting RIPK signaling may serve to limit the pathologic consequences of inflammatory astrocyte activation in PD and other neurodegenerative diseases.

Since the initial description of the A1 and A2 astrocyte subtypes, transcriptional profiling of astrocytes as an indication of their A1 or A2 activation state has grown increasing popular, though whether the respective transcriptomes of these states are stable across disease stimuli or reliably indicate neurotoxic vs. neurotrophic functional activity is controversial (12). In our study, treatment with α-synuclein PFFs induced neurotoxic activity and diminished phagocytic capacity, in line with previous descriptions of the A1 activation state (19, 58). However, while α-synuclein PFFs induced many putative A1-associated genes, we also saw marked upregulation of A2-associated genes as well, suggesting that the A1/A2 paradigm may not extend well, at least at the transcriptional level, to astrocyte activation downstream of fibrillar α-synuclein. One possible source of this discrepancy in our study was the use of human rather than murine astrocytes. While we analyzed expression of human orthologues of the A1- and A2-associated genes described in mice (19), it is possible that unique activation states in human astrocytes are marked by distinct transcriptional signatures that do not map directly onto those observed in rodents.

RIPK1 and RIPK3 are activators of necroptotic cell death (59, 60), which has been shown to have both protective and pathologic functions in diverse disease states, including infection (40, 61, 62), cancer (63), and sterile injury (64). In the CNS, RIPK signaling has principally been connected to deleterious neuroinflammation and necroptotic cell death in the context of neurodegenerative diseases, including PD (65, 66). However, we and others recently described a necroptosis-independent function for RIPK signaling in neurons in which a RIPK-dependent transcriptional program induced protective neuroinflammation and immunometabolic changes in response to neurotropic viral infection (34, 35). Here, we show that necroptosis-independent RIPK signaling can also coordinate inflammatory transcription in astrocytes, which is associated with the conferral of neurotoxic activity. These findings suggest that cell death-independent functions of this pathway in the CNS may extend beyond neurons, possibly to all cells of common neuroectodermal lineage.

Our study also identifies NF-κB as the central transcriptional mediator of astrocyte activation in response to α-synuclein PFFs. The activation of NF-κB in astrocytes has also previously been observed in a toxin-based model of PD (67). Notably, we found that α-synuclein PFFs both increased expression of NF-κB family members as well as induced nuclear translocation of p65, suggesting that this pathway may exhibit feed-forward regulation. We also show that RIPK signaling was required for NF-κB activation downstream of α-synuclein PFFs in astrocytes, in line with previous studies showing that RIPK signaling is a potent activator of NF-κB signaling (48-50, 68). However, previous work showed that RIPK-mediated transcriptional activation in neurons in the context of viral infection was independent of NF-κB (35), suggesting that the transcriptional machinery engaged by RIPK signaling in the CNS may be cell type- and stimulus-dependent.

While we have identified a previously unknown function for RIPK signaling and NF-κB in promoting a neurotoxic astrocyte activation state, several questions remain. For example, the mechanisms by which aggregated α-synuclein is sensed by astrocytes prior to the induction of inflammatory activation is poorly understood. While some studies have suggested that α-synuclein aggregates engage TLR4-mediated innate sensing pathways (24), whether adult astrocytes express TLR4 *in vivo* is a matter of debate (19, 69, 70). Further work, particularly in rodent models, will help further refine our understanding of the upstream signaling events that promote RIPK3 activation in astrocytes in the context of synucleinopathy. Finally, while several groups have now shown that activated astrocytes express some secreted factor that exerts neurotoxic activity, the identity of this factor (or factors) and its mechanism of action have been difficult to discern (11). Ongoing work characterizing the secretomes of activated astrocytes is needed to answer these important questions.

## Methods

### α-synuclein pre-formed fibrils

Purified human α-synuclein monomers were purchased from Proteos, Inc. (Kalamazoo, MI, #RP-003) and were used to generate PFFs according to established protocols (44). Briefly, monomers were diluted to a concentration of 5 mg/mL with PBS and agitated on a thermomixer at 1,000 RPM at 37°C for 7d. Aggregates were then diluted to a concentration of 100 µg/mL with 1x HBSS buffer followed by three 10s pulses of sonication using a 1/8” probe equipped QS5 Sonicator (Covaris, Woburn, MA). For cell culture experiments, freshly sonicated PFFs were used at 0.1 µg/mL.

### Inhibitors

BAY 11-7085 (#1743), SR 11302 (#2476), and Pyridone 6 (#6577) were purchased from Tocris Bioscience (Bristol, UK). JSH-23 was purchased from Selleck Chemicals (#S7351, Houston, TX). GSK963 (#SML2376), GSK872 (#530389), necrosulfanamide (#480073), and Z-VAD-FMK (#627610) were purchased from Millipore Sigma (Burlington, MA). All inhibitors were solubilized in DMSO. BAY 11-7085, SR 11302, and Pyridone 6 were used at a final concentration of 100 µM for cell culture treatments. JSH-23 was used at 50 µM. GSK 963 and GSK 872 were used at 1 µM. Necrosulfanamide was used at 10 µM. Z-VAD-FMK was used at 5 µM.

### Human astrocyte and neuronal cultures

Primary human midbrain astrocytes (#1850) were obtained from ScienCell Research Laboratories (Carlsbad, CA) and cultured in astrocyte media (AM, #1801), supplemented with 2% heat-inactivated fetal bovine serum (#0010), astrocyte growth supplement (#1852), and penicillin/streptomycin cocktail (# 0503). Cells were cultured in poly-L-lysine coated T75 flasks. Human neuronal cells SH-SY5Y (ATCC, Manassas, VA, #CRL-2266) were cultured in DMEM medium (VWR, Radnor, PA, #0101-0500) supplemented with 10% FBS (Gemini Biosciences West Sacramento, CA, #100-106), non-essential amino acids (Hyclone, #SH30138.01), HEPES (Hyclone #30237.01), penicillin, streptomycin, and antifungal (Gemini Biosciences #400-110, #100-104). SH-SY5Y cells were propagated in T75 flasks prior to the differentiation process. Cells were maintained in a humidified environment with 5% CO_2_ at 37°C.

### SH-SY5Y differentiation

SH-SY5Y neuroblastoma cells were differentiated into mature neuron-like cells by treating with retinoic acid (4 µg/mL; Sigma-Aldrich, St. Louis, MO, #R2625) and BDNF (25 ng/mL, Sigma-Aldrich, #B3795) diluted in DMEM supplemented with 2% heat inactivated fetal bovine serum (FBS, Gemini Biosciences, West Sacramento, CA, #100-106), non-essential amino acids (1x; HyClone, #SH30238.01), HEPES buffer (10 mM; HyClone, #SH30237.01), L-Glutamine, penicillin, streptomycin (Gemini Biosciences, #400-110), and antifungal amphotericin B (Gemini Biosciences, #100-104). Differentiated SH-SY5Y cultures were used for experiments 7d post-differentiation.

### Cell death and viability assay

Cell viability was assessed with the CellTiter-Glo Luminescent Cell Viability Assay kit (Promega, Madison, WI, #G7573), according to manufacturer’s instructions. Luminescence signal was read with a SpectraMax iD3 plate reader (Molecular Devices, San Jose, CA).

### Caspase 3/7 activity assay

Caspase 3/7 activity was measured using a chromogenic DEVD cleavage assay according to manufacturer’s instructions (R&D Systems, Minneapolis, MN, #K106-100).

### Quantitative real-time PCR

Total RNA from cultured cells were isolated with Qiagen RNeasy mini extraction kit (Qiagen, Valencia, CA, #74106) following the manufacturer’s protocol. RNA concentration was measured with a Quick Drop device (Molecular Devices, San Jose, CA). cDNA was subsequently synthesized with qScript cDNA Synthesis Kit (Quantabio, Beverly, MA, #95047). qRT-PCR was performed with SYBR Green Master Mix (Bio-Rad, Hercules, #CA1725125) using a QuantStudio5 instrument (Applied Biosystems, Foster City, CA). Cycle threshold (CT) values for analyzed genes were normalized to CT values of the housekeeping gene *18S* (CT_Target_ - CT_18S_ = ΔCT). Data were further normalized to baseline control values (ΔCT_experimental_ - ΔCT_control_ = ΔΔCT (DDCT). Primers were designed using Primer3 (https://bioinfo.ut.ee/primer3/) against human genomic sequences. A list of all primer sequences in our study appears in **Supplemental Table 1**.

### Immunocytochemistry

For imaging experiments, cells were grown on poly-D-lysine coated coverslips (Neuvitro, Vancouver, WA, #GG-12-PDL). Following experimental treatments, cells were fixed in 4% paraformaldehyde for 15m, followed by three washes in 1x PBS, followed by incubation in blocking solution (10% goat serum, Gibco, Waltham, MA, #16210 and 0.1% Triton X-100) for 30m at room temperature. Cells were then incubated in primary antibody (rabbit-anti-p65/RELA; 2 ug/mL; ThermoFisher, Waltham, MA, #10745-1-AP) diluted in blocking solution for 1h. After three 15m washes in 1x PBS, coverslips were incubated in secondary antibody (goat-anti-rabbit IgG conjugated Alexa Fluor 594; 2 ug/mL; Invitrogen Waltham, MA, #A32740) and nuclear stain (DAPI; 10 ug/mL; Biotium, Fremont, CA, #40043) for 15m at room temperature, followed by another series of washes with 1x PBS. Coverslips were than mounted using ProLong Diamond Anitfade Mountant (Invitrogen, #P36931) onto slides. Images were acquired with an Airyscan fluorescent confocal microscope (Carl Zeiss LSM 800).

### TUNEL assay

TUNEL was performed using a standard kit according to the manufacturer’s protocol (TMR In Situ Cell Death Detection Kit, Sigma-Aldrich, #12156792910) in combination with nuclear staining using DAPI (10 ug/mL; Biotium, Fremont, CA, #40043). Images were captured using a 20x objective. Numbers of TUNEL positive nuclei in each image were counted by a blinded operator.

### Colocalization analysis

For each coverslip, three regions with matched cell density were captured with the 63x objective. p65 colocalization with DAPI was quantified using the Colocalization Colormap plugin in ImageJ software (National Institute of Health, Bethesda, MD). The plugin calculates normalized mean deviation product, an index of correlation between pixels. Fisher’s Z transformation was applied to the index of correlation prior to comparison.

### Nuclear protein extraction

Primary human midbrain astrocytes were cultured to confluency. Cells were washed twice with PBS followed by 5m incubation in cold 5 mM EDTA. Cells were then scraped into 15 mL conical tubes and centrifuged for 5m at 1,000 rpm. Nuclear extraction was performed using a standard kit according to the manufacturer’s protocol (Nuclear Extraction Kit, abcam, Cambridge, MA, #ab113474).

### NF-κB p65 transcription factor assay

Protein concentrations of nuclear extracts were determined using BCA assay (ThermoFisher, #23227), according to manufacturer’s instructions. Equal amounts of protein were then processed through an ELISA based kit for detecting p65 (NF-κB p65 Transcription Factor Assay Kit, abcam, Cambridge, MA, #ab133112). Absorbance at 450 nm was read with SpectraMax iD3 plate reader (Molecular Devices, San Jose, CA).

### Statistical analysis

Data analysis for most studies was performed with two-way analysis of variance (ANOVA) with Sidak’s correction for multiple comparisons using GraphPad Prism Software v8 (GraphPad Software, San Diego, CA). P<0.05 was considered statistically significant. Analysis of publicly available microarray data was performed in GEO2R and the GO Enrichment Analysis tool (71). Corrected p-values (false discovery rate) were determined using the Benjamini & Hochberg procedure.

## Acknowledgements

This work was supported by a research grant from the American Parkinson’s Disease Association and startup funds from Rutgers University (to BPD). MK was supported in part by a Parkinson’s Foundation Summer Student Fellowship. APP was supported in part by a Division of Life Sciences Summer Undergraduate Research Fellowship from Rutgers University.

## Ethics declarations

The authors declare they have no conflicts of interest.

