## Supplemental Material for "Fibrillar α-synuclein induces neurotoxic astrocyte activation via RIP kinase signaling and NF-κB"

### Supplemental Data

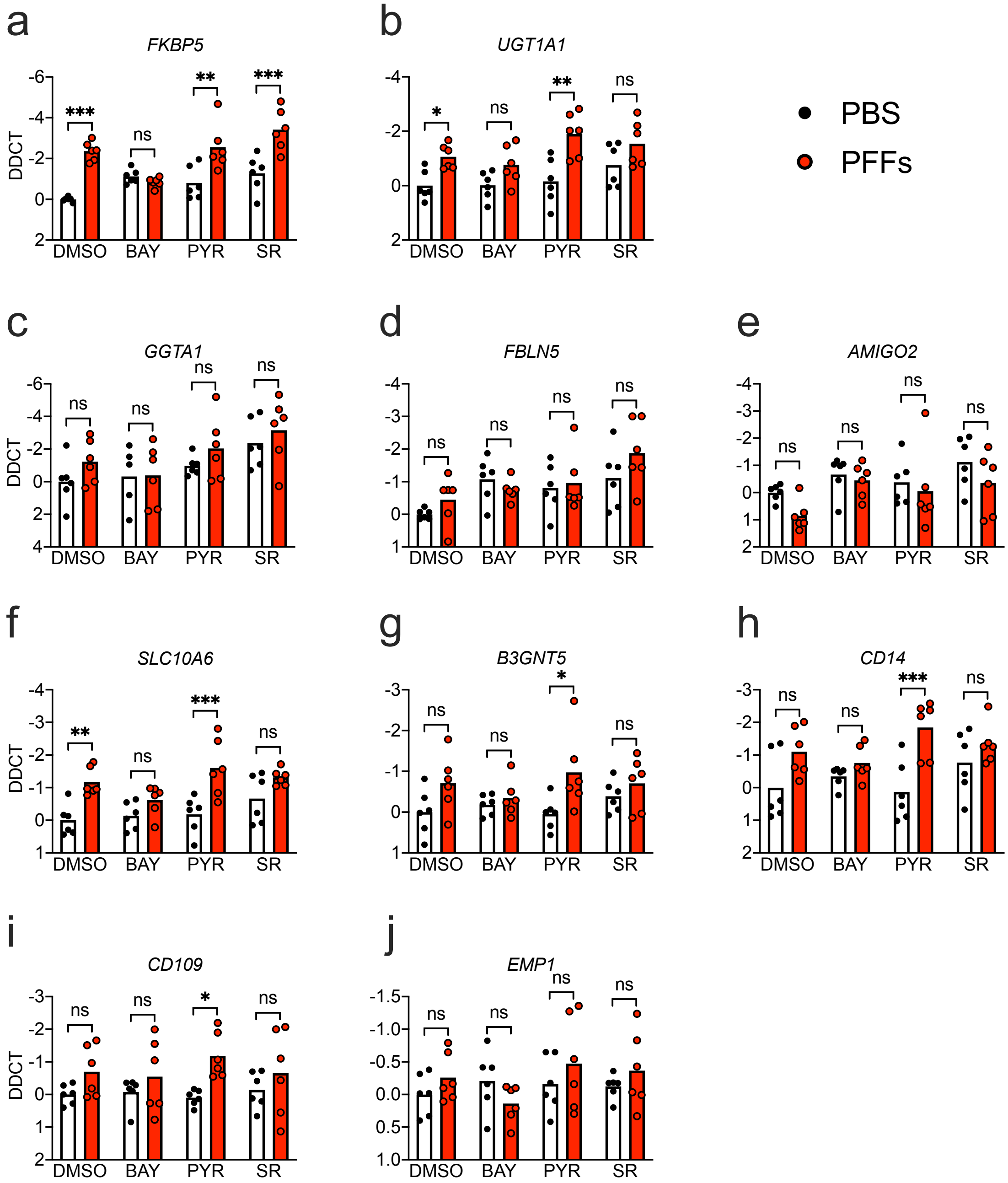


**Supplemental Figure 1. α-synuclein PFFs induce NF-κB-dependent transcriptional activation in human midbrain astrocytes.**

**a-j)** Primary human midbrain astrocyte cultures were treated for 24h with PFFs or PBS control solution. Cultures were pretreated (30min) with inhibitors of NF-κB (BAY), JAK/STAT (PYR), or AP1 (SR) signaling prior to addition of PFFs**.** Levels of indicated A1-associated transcripts **(a-e)** and A2-associated transcripts **(f-j)** were measured using qRT-PCR. ns: not significant, *$p < 0.05$, **$p < 0.01$, ***$p < 0.001$. Bars represent group means. n=6 independent replicates for all experiments.

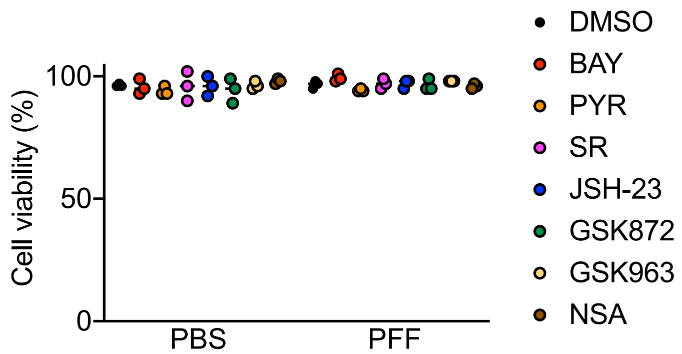


**Supplemental Figure 2. Transcription factor and necroptosis pathway inhibitors are not toxic to differentiated SH-SY5Y cultures.**

Differentiated cultures of SH-SY5Y cultures were treated with the indicated inhibitors for 24h. Cell viability was then assessed using an ATP-luciferase assay (CellTiter Glo). n= 3 independent replicates.

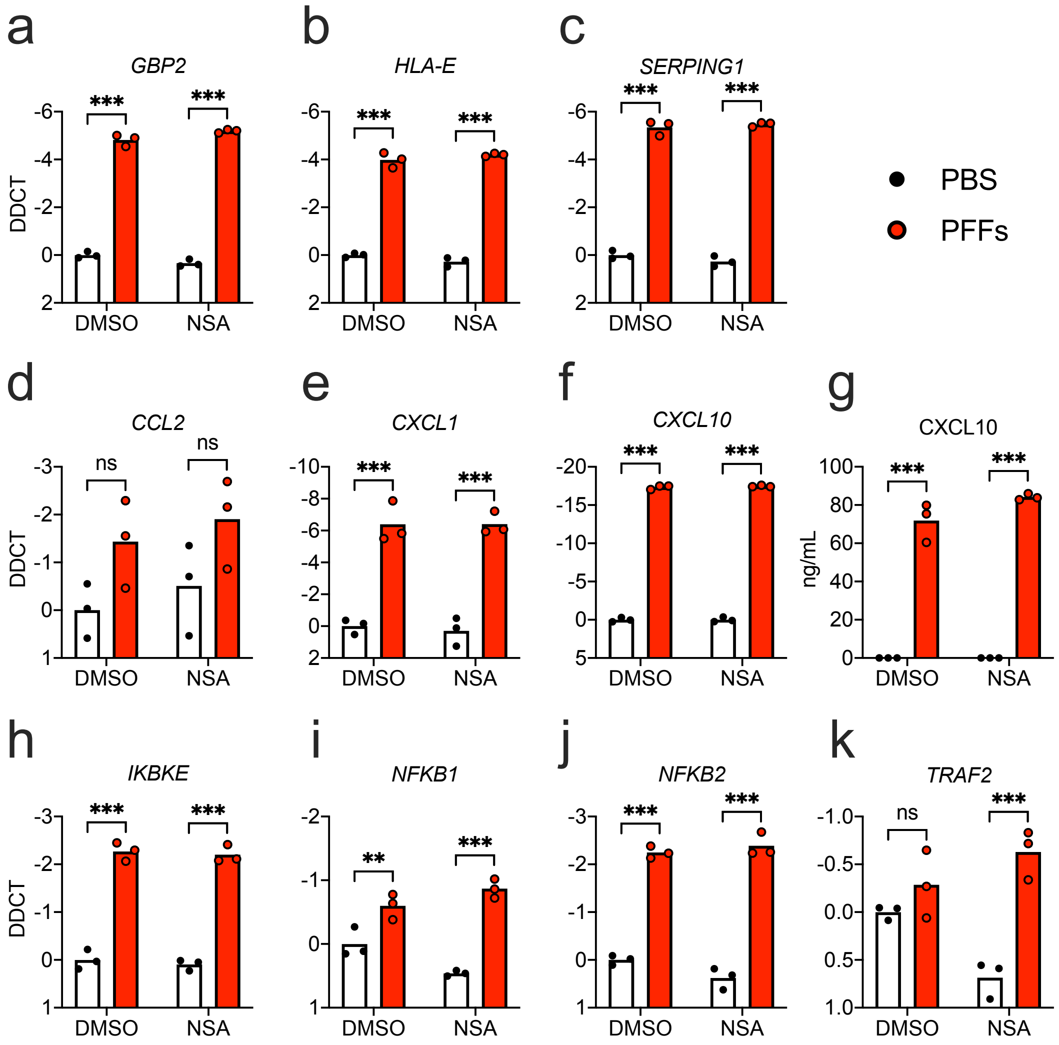


**Supplemental Figure 3. α-synuclein PFF-mediated transcriptional activation in astrocytes does not require MLKL**

**a-j)** Primary human midbrain astrocyte cultures were treated for 24h with PFFs or PBS control solution. Cultures were pretreated (30min) with MLKL inhibitor (NSA) prior to addition of PFFs**. a-f, h-k)** Levels of indicated transcripts were measured using qRT-PCR. **g)** Levels of CXCL10 protein in culture supernatants were measured via ELISA. ns: not significant, **p < 0.01, ***p < 0.001. Bars represent group means. n=3 independent replicates for all experiments.

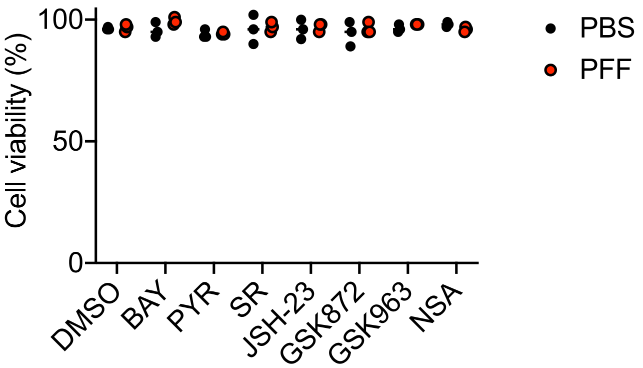


**Supplemental Figure 4. Neither PFFs nor transcription factor/necroptosis pathway inhibitors induce cell death in human midbrain astrocyte cultures.**
Primary human midbrain astrocyte cultures were treated with PFFs or PBS along with indicated inhibitors for 24h. Cell viability was then assessed using an ATP-luciferase assay (CellTiter Glo). n= 3 independent replicates.

| Gene | Forward | Reverse |
|---|---|---|
| *18S* | AGAAACGGCTACCACATCCA | CCCTCCAATGGATCCTCGTT |
| *AMIGO2* | CTTCAGCGTTTGGAGGGCT | CAGGGAACAGTCACAGACAAAT |
| *AXL* | CCAGGACACCCCAGAGGTGCTAAT | TGGTGGACTGGCTGTGCTTGC |
| *B3GNT5* | ACTCCTCCCCAACAAGGTCT | TTTAACCCCAAACTGGCAAC |
| *CCL2* | GCAGCAAGTGTCCCAAAGAA | CTGGGGAAAGCTAGGGGAAA |
| *CD109* | CAGGAATGTGGACTCTGGGT | CTTTCGGACATGTGGACTGC |
| *CD14* | CCGCTGTGTAGGAAAGAAGC | GCAGCGGAAATCTTCATCGT |
| *CLCF1* | GCACAGAGTGGCAAACAAAA | ACACCCCAAAATGCTACTGC |
| *CXCL1* | ACTCTACCTGCACACTGTCC | TCCCCTGCCTTCACAATGAT |
| *CXCL10* | GTGGCATTCAAGGAGTACCTC | TGATGGCCTTCGATTCTGGATT |
| *EMP1* | CCAGTACACCAGCAGAGGAA | AACAGTAGCGATGTGGACCA |
| *FBLN5* | TCGCCAGTCAGGACAGTGT | AGTAGGGGTTCGAGTAGGGC |
| *FKBP5* | CTCCCTAAAATTCCCTCGAATGC | CCCTCTCCTTTCCGTTTGGTT |
| *GAS6* | ATCAAGGTCAACAGGGATGC | CTTCTCCGTTCAGCCAGTTC |
| *GBP2* | CTATCTGCAATTACGCAGCCT | TGTTCTGGCTTCTTGGGATGA |
| *GGTA1* | ATGACAGCAGTGCTCAGAAGG | AGCCGAAGCTCTGTTGTGT |
| *HLA-A* | GACCAGGAGACACGGAATGTG | CCTCGTTCAAGGCGATGTAATC |
| *HLA-E* | TTCCGAGTGAATCTGCGGAC | GTCGTAGGCGAACTGTTCATAC |
| *IKBKE* | TGCGTGCAGAAGTATCAAGC | TACAGGCAGCCACAGAACAG |
| *LCN2* | GAAGTGTGACTACTGGATCAGGA | ACCACTCGGACGAGGTAACT |
| *MEGF10* | TGACTGCTTGCCTGGCTTCACA | GTTACAGGTTCCGTTGTTGGTGC |
| *MERTK* | CAGGAAGATGGGACCTCTCTGA | GGCTGAAGTCTTTCATGCACGC |
| *NFKB1* | GCAGCACTACTTCTTGACCACC | TCTGCTCCTGAGCATTGACGTC |
| *NFKB2* | GGCAGACCAGTGTCATTGAGCA | CAGCAGAAAGCTCACCACACTC |
| *PSMB8* | GGTCCTACATTAGTGCCTTACGG | CGCAGATAGTACAGCCTGCATT |
| *PTGS2* | TGAGCATCTACGGTTTGCTG | TGCTTGTCTGGAACAACTGC |
| *PTX3* | GTGGGTGGAGAGGAGAACAA | TTCCTCCCTCAGGAACAATG |
| *S100A10* | ATGAAGGACCTGGACCAGTG | GCAGATTCCTTAAGCGACCC |
| *SERPING1* | GGGATGCTTTGGTAGATTTCTCC | GAGGATGCTCTCCAGGTTTGT |
| *SLC10A6* | TGTTGCCGATGTTCATTTGT | CACTGTGAAAACGAGCTCCA |
| *SPHK1* | ACCCATGAACCTGCTGTCTC | CAGGTGTCTTGGAACCCACT |
| *SRGN* | GGACTACTCTGGATCAGGCTT | CAAGAGACCTAAGGTTGTCATGG |
| *TRAF2* | CACCGGTACTGCTCCTTCTG | TGAACACAGGCAGCACAGTT |
| *UGT1A1* | TGTCCCATGCTGGGAAGATAC | GAATGGCACAGGGTACGTCT |

**Supplemental Table 1: Primer sequences for qRT-PCR**
